# Twenty-four hour face mask sampling in pulmonary tuberculosis reveals three distinct patterns of bacterial aerosol production dissociated from conventional markers of transmission risk

**DOI:** 10.1101/426825

**Authors:** CML Williams, M Abdulwhhab, SS Birring, E De Kock, NJ Garton, A Stoltz, P Haldar, MR Barer

## Abstract

**Rationale:** Although tuberculosis (TB) is transmitted by *Mycobacterium tuberculosis* (Mtb) in aerosols, little is known of the dynamic characteristics of spontaneous output of bacilli in this form. We have developed and implemented a mask aerosol sampling system (MASS) for longitudinal capture and study of spontaneous aerosol.

**Objective:** To determine patterns of Mtb output in aerosols, captured using the MASS over 24 hours and their association with existing criteria used to assess transmission risk in patients with pulmonary TB.

**Methods:** Twenty-four hospitalised patients with newly diagnosed pulmonary TB recruited in Pretoria, South Africa, wore FFP1 masks for one hour out of every three for 24 hours. Aerosol was captured in a gelatine filter processed for Mtb quantitation by PCR. Serial sputum was collected and objective cough frequency monitoring performed over the same period.

**Measurements and Main Results:** Mtb was detected in 86.5% of 192 mask samples and 20.7% of 38 assessable sputum samples obtained from the cohort. Mtb was detected by MASS in all but two patients. Three dynamic patterns of expression were identifiable in Mtb aerosol producers: i. variable high; ii. consistent; and iii. variable low. No diurnal variation was apparent and there was no correlation between mask Mtb and either sputum Mtb levels or cough frequency. Sputum smear status, culture time to positivity and chest radiographic characteristics also failed to associate with MASS bacillary output.

**Conclusions:** Conventional markers of tuberculosis case infectivity do not predict bacillary aerosols detected by the MASS. The MASS provides a novel, non-invasive tool for tuberculosis diagnosis and control.

## Introduction

Community and patient level control measures of Tuberculosis (TB) currently rely on clinical surrogates of infectivity. These include presence and frequency of cough, extent of cavitatory disease on chest radiograph (CXR), sputum smear acid-fast bacillary (AFB) or Xpert MTB/RIF grade and bacterial burden in culture^1^.

These indicators are chosen on the basis that they reliably inform the risk of onward transmission of the infection by patient generated aerosols from those individuals with marked pulmonary disease. However, the overall contribution of this patient group to transmission across populations is uncertain. It would clearly be desirable to replace these indirect markers of infectivity with direct assessment of patient generated aerosols, both in specific cases assessment and in broader programmes of active case finding.

Evidence for the airborne transmission of TB was first presented by Riley and colleagues (ref) using a guinea pig model, where animals are exposed to the room air from pulmonary TB patients. Recently, Fennelly and colleagues have captured culturable infectious aerosols containing *Mycobacterium tuberculosis* (Mtb) directly from patients using the Cough Aerosol Sampling System (CASS)^2^. While the Riley model has not been directly related to community level transmission by individuals, CASS based studies have provided early reports in this regard and analyses have shown aerosol is superior to sputum smear grade as a measure of transmission risk^3^. Neither of these approaches examine the characteristics of aerosol production by individuals over time. Infectivity measured by the guinea pig model is not time resolved beyond cumulative exposure and CASS is based on a single episode of forced coughing that may not reflect natural aerosol production.

We have previously reported on a novel Mask Aerosol Sampling System (MASS) that captures Mtb directly from patients under natural conditions^4^, offering the possibility for studying this important question.

In this prospective study, we have utilised MASS to investigate the dynamic patterns of Mtb output, including possible diurnal variation, in patient-generated aerosols over 24 hours. We compare this with Mtb in expectorated sputum over the same period and investigate the association of these direct measures of infectivity with conventional measures of transmission risk commonly used in clinical practice. Our key hypotheses are that sputum bacillary content, cough frequency and extent disease on CXR are associated with Mtb detected in mask samples.

## Methods

### Study Population

Hospitalised pulmonary TB patients diagnosed by either Xpert MTB/RIF or smear AFB positive on a sputum sample were invited to take part at three hospital sites in Pretoria, South Africa (Steve Biko Academic Hospital, Tshwane District Hospital or Kalafong District Hospital) between September and December 2015. Participants were eligible if they were 18 years or older, did not require continuous oxygen therapy by face mask or nasal prongs and were untreated or within 24 hours of starting anti-TB treatment.

### Ethical consideration

Patients were consented in their preferred language and provided written consent. This study was approved by the Faculty of Health Sciences Research Ethics Committee at the University of Pretoria and Provincial approval from Gauteng Health Department (ref: RET_215_UP03).

### Aerosol and Sputum Sampling

Each participant wore a modified FPP1 face mask containing a gelatine filter (pore size 0.3μm, Sartorius, Germany). They were given no set instructions on vocal manoeuvres during this time and allowed to cough, talk, laugh or sleep as desired. If they needed to expectorate then they were asked to lift the mask briefly, after coughing, to expectorate into a sputum collection pot.

Each participant underwent MASS for an hour every three hours and were provided with a new sputum pot at every three hour interval and encouraged to collect whatever they expectorated spontaneously for the 24 hour study duration.

They were observed to ensure that the mask was worn for the whole hour of sampling and participant behaviour was recorded, including anytime the mask was directly in front of the mouth. Sleep was documented if the participant had their eyes closed for >10 minutes and were not obviously rousable by noise, resting was noted if the participant had their eyes closed for <10 minutes at a time and/ or had their eyes open but not engaging in any activity such as talking, eating, reading etc. Other activities such as eating, washing, reading and talking were documented.

To provide background controls swabs were taken from a total of 72/192 masks, holders and transport bags to address the possibility of environmental contamination under the same conditions as the processed masks. All were negative by PCR.

### Initial processing of Aerosol and Sputum samples

Gelatine filters were dissolved in 1.5mls of 2% w/v NaOH and incubated at room temperature for 15 minutes before neutralising with 190μl 4M HCL. Samples were agitated by hand at 0 and 8 minutes. The dissolved filter was then centrifuged at 13,400 *xg* for 10 minutes, the supernatant removed and the pellet overlaid with 100μl of TE buffer (20mM Tris and 2mM EDTA pH 8.0), prior to storage at - 80°C.

As in our previous work^4^, we have assigned bacillary output recovered from this mask sample to be in aerosols, although we acknowledge that a range of particles including those outside the respirable range are likely included. We have used the terms mask and aerosol sampling interchangeably to denote results from this sample.

Sputum was decontaminated and processed in accordance with Turapov and colleagues^5^, before being re-suspended in 1ml of the supernatant, passaged through a 23G blunt needle 10 times and centrifuged at 13,400×g for 10 minutes. The supernatant was removed and the pellet overlaid with 100μl of TE buffer (20mM Tris and 2mM EDTA pH 8.0) prior to storing at -80°C.

### Extraction and quantification of DNA from bacterial pellets

Bacterial pellets from the aerosol and sputum samples underwent the same in-house extraction method modified from that outlined by Reddy and colleagues^6^. 100μl Chelex-NP40 (50% w/v Chelex-100, 1% w/v Non-idet P40, 1% w/v Tween 20) was added to the defrosted bacterial pellet along with 0.3g glass beads (150-212μm Sigma-Aldrich USA). The sample was then homogenised in a Fast Prep at 6.5 m/s for 45 seconds and ice incubated for 5 minutes. This homogenisation step was repeated 3 further times prior centrifugation at 13,400×g for 2 minutes. 200μl of the supernatant was heated at 95°C for 30 minutes prior to removal from the CL3.

Copy numbers of IS6110 were quantified in each sample by real-time q-PCR run on a Rotor-Gene (Qiagen UK) using a TaqMan IS6110 assay outlined by Akkerman and colleagues^7^. Real-time PCR signals were analysed by Rotor-Gene 6000 Series Software 1. The functions “slope correct” and “ignore cycles” were applied to analyses. In accordance with Dorak and colleagues only runs with correlation coefficients (R^2^) and reaction efficiencies above 0.99 and 0.8 respectively were included for analysis^8^. Technical replicates were undertaken in triplicate and considered reproducible if the difference in cycle threshold (Ct) was less than 1.

As differences in IS6110 copy number between bacterial strains limits confidence in comparative analysis, a subset of samples for each participant was further analysed using the Mtb-specific RD9 DNA sequence, single copy gene^9^. Comparative statistical analyses have been assessed using both IS6110 copy numbers and genome copy number derived using the ratio between IS6110 and RD9 (see supplemental data).

### Cough Sampling

A Leicester Cough Monitor (LCM) was worn by each participant for the 24 hours of the study. As described by Birring and colleagues, the LCM consists of an MP3 recorder (Sony ICD PX333) worn at the participant’s waist in protective bag connected to a clip microphone (Philips LFH9165) that is worn as close to the sternoclavicular joint as possible^10^. Recordings are analysed using specialised semi-automated software as previously described, that anonymised sounds and quantified coughs both as single events or cough bouts^10^. The LCM position was checked every 3 hours throughout the study. As well as being well validated for use in patients with many respiratory conditions including TB^10-13^the LCM was validated for use with MASS prior to this study.

A subset of the recordings were validated by a second member of the research team, who was ‘blinded’ to the patient information and the primary analysis of the recording. There was good agreement in the subset reviewed by both members of the research team with an Intra-Class Correlation of R=0.99 (CI 0.996-1.00).

Nocturnal and daytime cough were defined by the time periods of 23:00-05:00 and 05:00-23:00, respectively. These periods were determined by observations of both the participants and working patterns of the hospital ward.

### Participant Data Collection

Clinical and demographic data was collected for each participant including routine microbiological investigations; sputum AFB smear microscopy, Xpert MTB/RIF and liquid culture (BACTEC MGIT 960). Radiological changes were graded using methods described by Ralph and colleagues^14^. Duration of symptoms was determined by questionnaire.

### Data Analysis

Data was analysed using GraphPad prism (Version 7 for Windows, USA) Excel (2010 for Windows Microsoft USA), and SPSS Statistics (Version 22, IBN, USA). The distribution of data was assessed using the Kolmogorov-Smirnov test and found to be non-normally distributed. Therefore averages of groups were reported Median (IQR), comparisons were carried out using the Mann–Whitney U-test or Wilcoxon signed rank test and correlations between variables were analysed with Spearman’s correlation coefficient (ρ). A p-value <0.05 was considered statistically significant. Cohen’s Kappa coefficient test was used to assess agreement between aerosol and sputum results. Total Aerosol Mtb output was calculated by multiplying the sum of the IS6110 copy number detected in the 8 aerosol samples collected by 3.

## Results

We first established the quantitative features of our sampling system in vitro. Direct contamination of mask filters with Mtb dilutions of mid-exponential culture, revealed a limit of detection of 152 colony forming units. We further determined that exposure to aerosols of *M. bovis BCG* for 15 minutes confirmed the capacity of the system to detect a suitable dynamic range. (see supplemental data)

### Patient enrolment and clinical characteristics

Of 78 pulmonary TB patients screened in Pretoria, South Africa, 33 patients were ineligible and 20 declined participation either prior to or after study enrolment (Figure 1). One further patient was excluded from analysis after sputum sampling identified *M. intracellulare* infection (figure 1). For 16 of the 24 participants completing the study, all samples were taken prior to commencing TB treatment, with the other 8 receiving a single dose either prior to or during their respective sampling periods (Table 1).

**Table 1:**
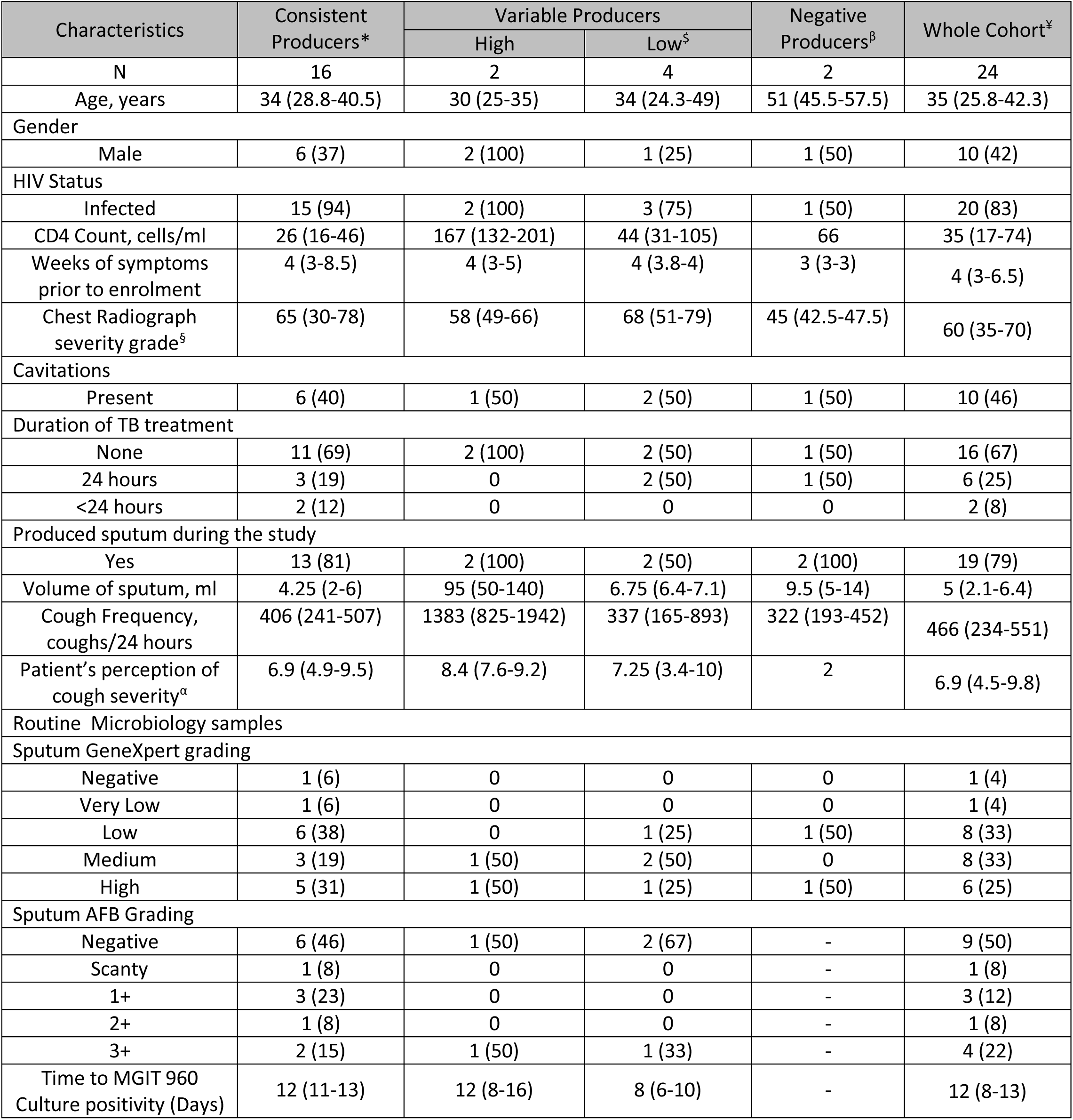
Characteristics of 24 patients with pulmonary TB studied, based on patterns of aerosol production. Values are median (IQR) or n (%) unless otherwise stated. ^§^Chest Radiograph severity grade based on extent of disease and presence of cavitation - range 0-140 (ref). ^α^Measured using Visual Annalogue Score (VAS) range 1-10. Definition of abbreviations: IQR= Interquartile range, AFB= Acid Fast Bacilli. *Missing data for consistent aerosol producers: Sputum AFB grade (3), MGIT 960 Culture (3), VAS (1), Chest Radiography severity grade and Cavitations (1) ^$^Missing data for Low Variable producers: Sputum AFB grade (1) & MGIT 960 Culture (2) ^β^Missing data for Negative producers: Sputum AFB grade (2), MGIT 960 Culture (2), VAS (1) ^¥^Missing data for Whole cohort: Sputum AFB grade (6), MGIT 960 Culture (7), VAS (2), Chest Radiography severity grade and Cavitations (1)

| Characteristics | Consistent Producers* | Variable Producers |  | Negative Producers<sup>β</sup> | Whole Cohort<sup>γ</sup> |
| --- | --- | --- | --- | --- | --- |
|  |  | High | Low<sup>§</sup> |  |  |
| N | 16 | 2 | 4 | 2 | 24 |
| Age, years | 34 (28.8-40.5) | 30 (25-35) | 34 (24.3-49) | 51 (45.5-57.5) | 35 (25.8-42.3) |
| Gender |  |  |  |  |  |
| Male | 6 (37) | 2 (100) | 1 (25) | 1 (50) | 10 (42) |
| HIV Status |  |  |  |  |  |
| Infected | 15 (94) | 2 (100) | 3 (75) | 1 (50) | 20 (83) |
| CD4 Count, cells/ml | 26 (16-46) | 167 (132-201) | 44 (31-105) | 66 | 35 (17-74) |
| Weeks of symptoms prior to enrolment | 4 (3-8.5) | 4 (3-5) | 4 (3.8-4) | 3 (3-3) | 4 (3-6.5) |
| Chest Radiograph severity grade<sup>§</sup> | 65 (30-78) | 58 (49-66) | 68 (51-79) | 45 (42.5-47.5) | 60 (35-70) |
| Cavitations |  |  |  |  |  |
| Present | 6 (40) | 1 (50) | 2 (50) | 1 (50) | 10 (46) |
| Duration of TB treatment |  |  |  |  |  |
| None | 11 (69) | 2 (100) | 2 (50) | 1 (50) | 16 (67) |
| 24 hours | 3 (19) | 0 | 2 (50) | 1 (50) | 6 (25) |
| <24 hours | 2 (12) | 0 | 0 | 0 | 2 (8) |
| Produced sputum during the study |  |  |  |  |  |
| Yes | 13 (81) | 2 (100) | 2 (50) | 2 (100) | 19 (79) |
| Volume of sputum, ml | 4.25 (2-6) | 95 (50-140) | 6.75 (6.4-7.1) | 9.5 (5-14) | 5 (2.1-6.4) |
| Cough Frequency, coughs/24 hours | 406 (241-507) | 1383 (825-1942) | 337 (165-893) | 322 (193-452) | 466 (234-551) |
| Patient's perception of cough severity<sup>α</sup> | 6.9 (4.9-9.5) | 8.4 (7.6-9.2) | 7.25 (3.4-10) | 2 | 6.9 (4.5-9.8) |
| Routine Microbiology samples |  |  |  |  |  |
| Sputum GeneXpert grading |  |  |  |  |  |
| Negative | 1 (6) | 0 | 0 | 0 | 1 (4) |
| Very Low | 1 (6) | 0 | 0 | 0 | 1 (4) |
| Low | 6 (38) | 0 | 1 (25) | 1 (50) | 8 (33) |
| Medium | 3 (19) | 1 (50) | 2 (50) | 0 | 8 (33) |
| High | 5 (31) | 1 (50) | 1 (25) | 1 (50) | 6 (25) |
| Sputum AFB Grading |  |  |  |  |  |
| Negative | 6 (46) | 1 (50) | 2 (67) | - | 9 (50) |
| Scanty | 1 (8) | 0 | 0 | - | 1 (8) |
| 1+ | 3 (23) | 0 | 0 | - | 3 (12) |
| 2+ | 1 (8) | 0 | 0 | - | 1 (8) |
| 3+ | 2 (15) | 1 (50) | 1 (33) | - | 4 (22) |
| Time to MGIT 960 Culture positivity (Days) | 12 (11-13) | 12 (8-16) | 8 (6-10) | - | 12 (8-13) |

**Figure 1.**
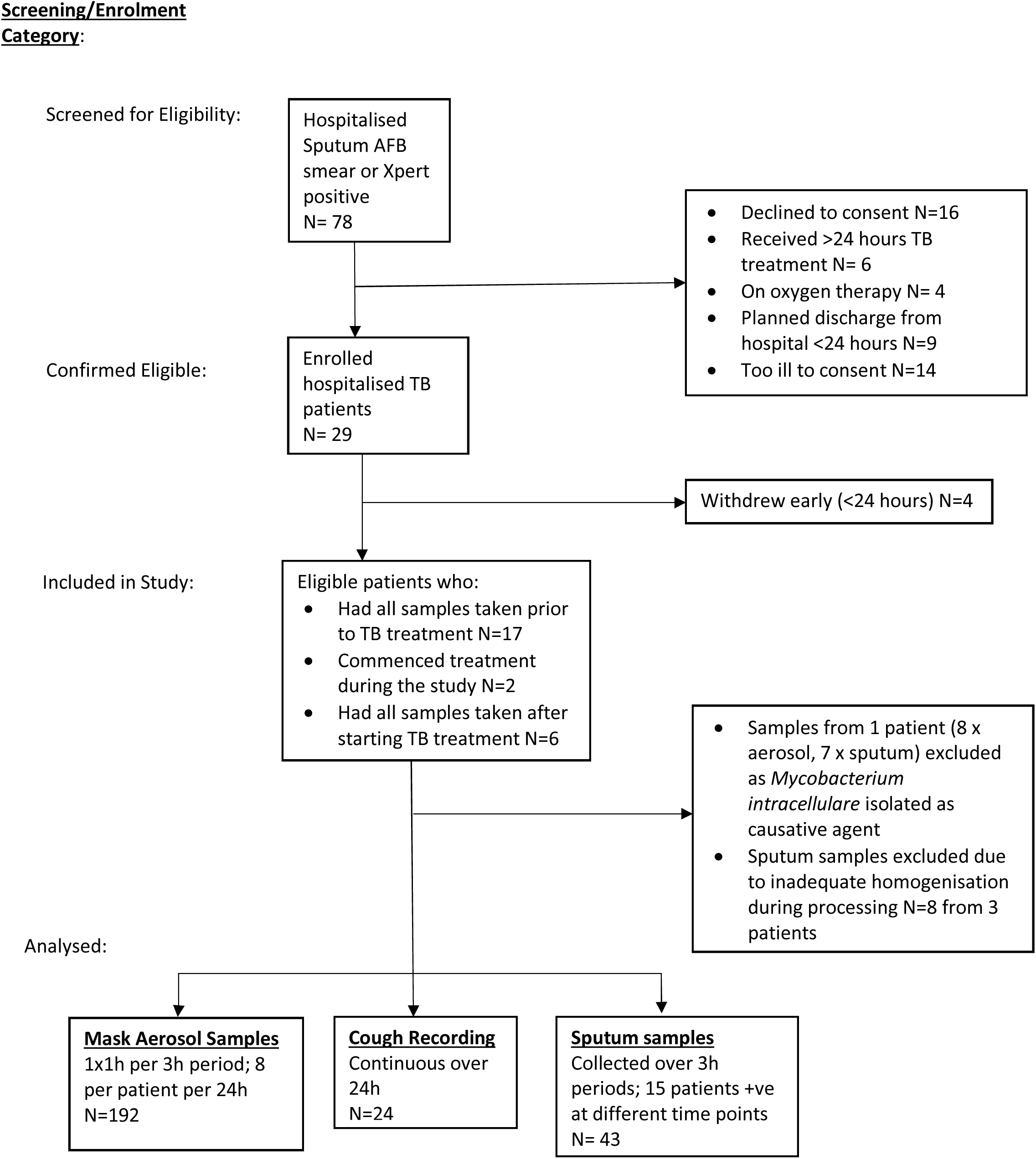
Study profile. AFB – acid-fast bacilli; TB - tuberculosis

All study participants were Black African and hospitalised with microbiologically confirmed advanced pulmonary TB (Table 1). The median age of the cohort was 35 years and 14 participants (58%) were female. HIV co-infection was identified in 20 participants (83%), with a median CD4^+^ cell count of 35 cells/ml, of whom 4 (20%) were receiving antiretroviral therapy at enrolment.

### Four patterns of aerosol production were observed

A one hour mask sample was readily obtained from all 24 participants during each of 192, three hour sampling periods. As evident in Figure 2, Mtb was detected by PCR in aerosol in all but two participants, with 17 participant’s positive in every mask. The dynamic pattern of Mtb output broadly fell into three groups over 24 hours we describe as consistent, variable high and variable low. For the majority of aerosol producers (16/22) the extent of variation was within 40 fold of their minimum detected hourly output, whilst the majority showed variation below 5 fold (see supplemental data); the remaining 6 participants produced several samples exceeding this limit. On this basis we determined four patterns of aerosol production, with respect to Mtb detection, in our cohort: i. Negative aerosol producers; ii. Consistent aerosol producers; iii. Low variable aerosol producers and iv. High variable aerosol producers (fig 2). These patterns were not influenced by the diurnal cycle.

**Figure 2:**
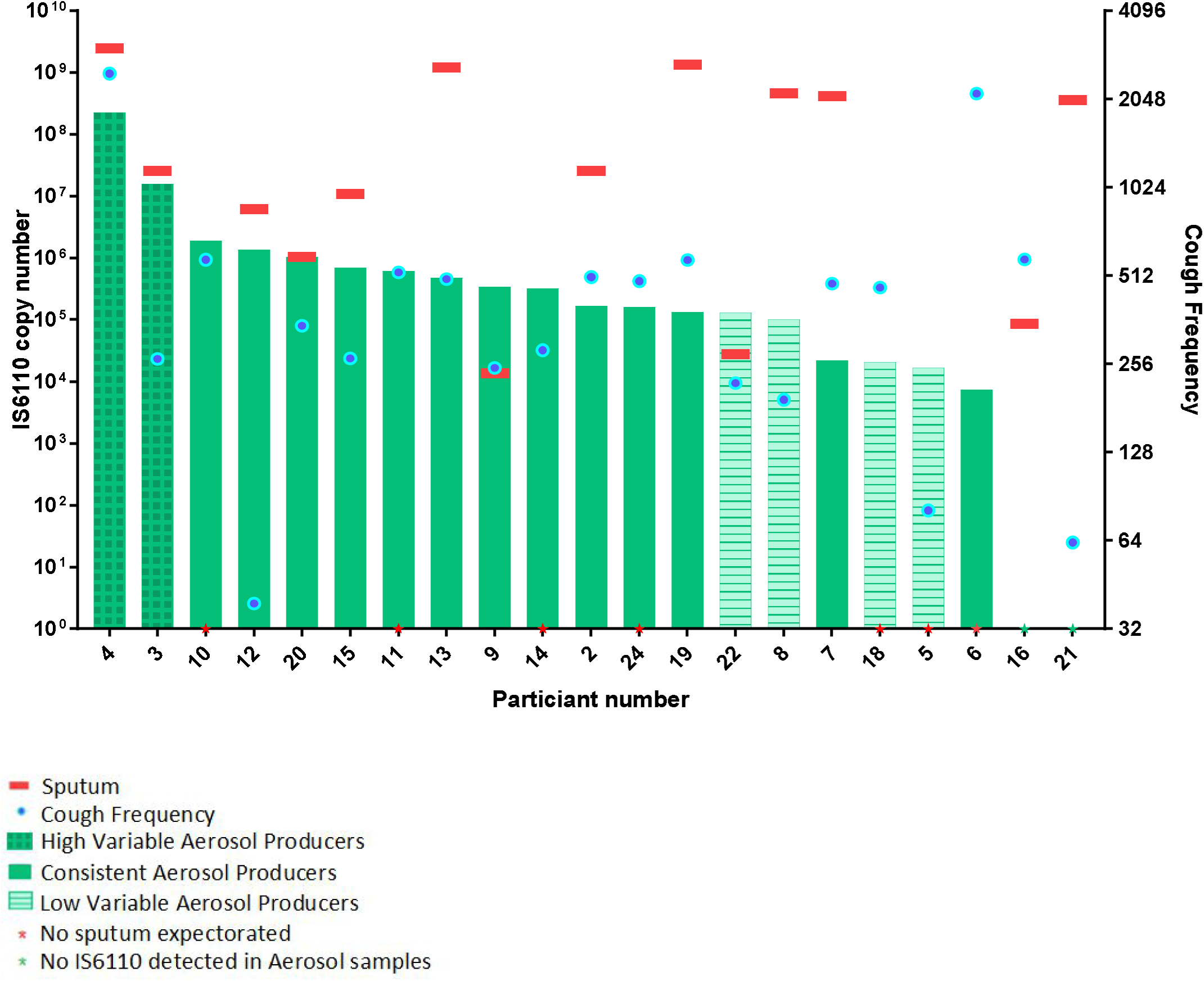
Total sputum and aerosol Mtb output, together with cough count over 24 hours, ranked by aerosol production in those with adequate aerosol and sputum sampling and processing.

**Figure 3:**
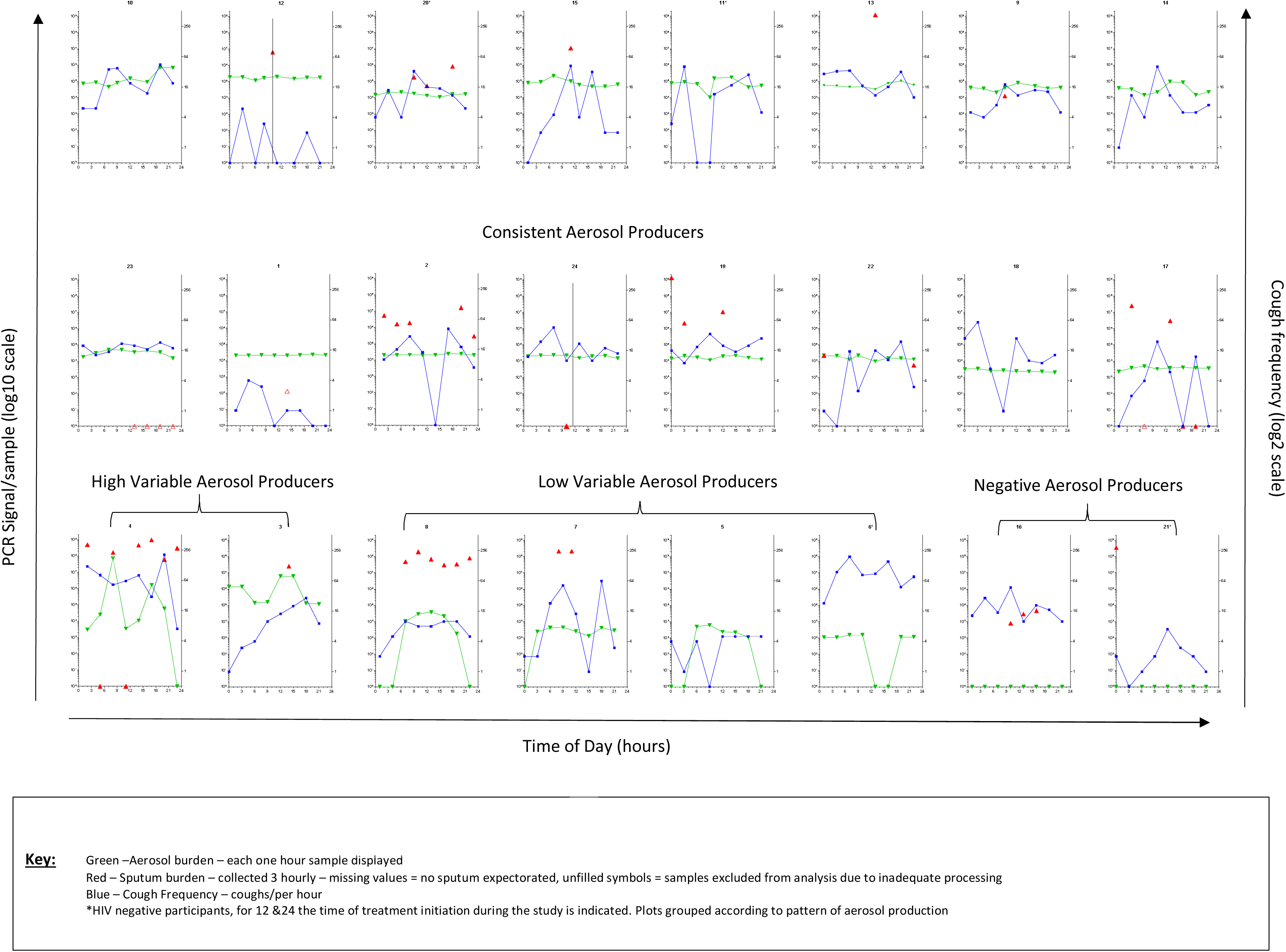
Pattern of 24 Hour Mycobacterial release by TB patients in aerosol and sputum compared with cough. **<u>Key:</u>** Green –Aerosol burden – each one hour sample displayed Red – Sputum burden – collected 3 hourly – missing values = no sputum expectorated, unfilled symbols = samples excluded from analysis due to inadequate processing Blue – Cough Frequency – coughs/per hour *HIV negative participants, for 12 &24 the time of treatment initiation during the study is indicated. Plots grouped according to pattern of aerosol production

Aerosol samples were taken at 68 time points when participants were observed to be sleeping for the full one hour sampling from 23/24 participants (with the exception of a brief rousing to place the mask on and off). Mtb was detected in 53 samples from 19 participants. The median (IQR) Mtb burden detected during sleeping was 1.4×10^4^ (5.0 × 10^3^ − 2.6×10^4^). In contrast, cough frequency was significantly lower during these sampling periods (median (IQR) 4 (1-17)). It was particularly striking that for 11 of these aerosol samples Mtb was detected with no occurrence of cough (fig 4).

**Figure 4:**
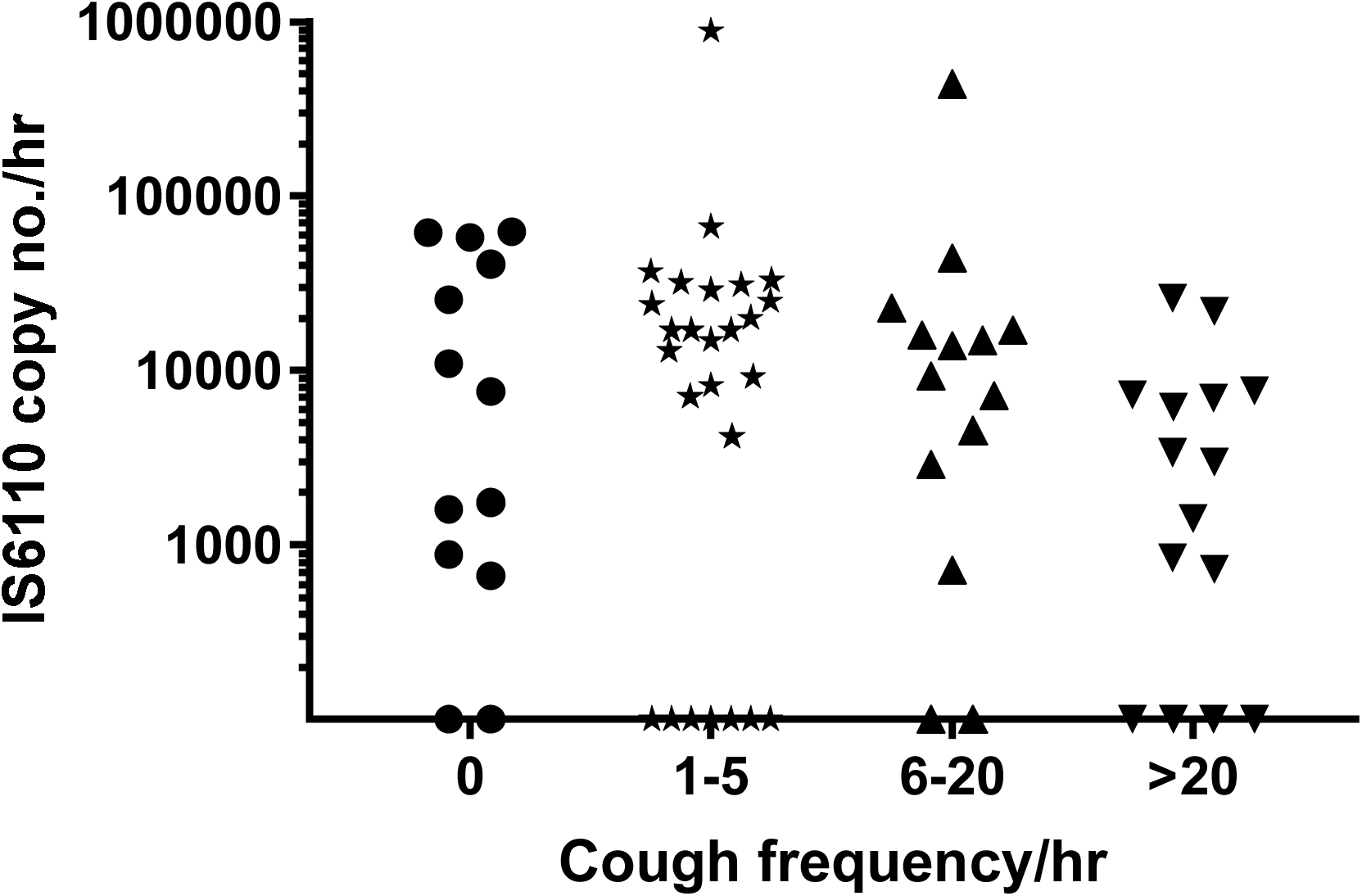
Aerosol Mtb burden detected in samples taken when participants were sleeping. grouped by cough recorded during sampling. Note: Mtb levels in samples plotted on x axis were below the limit of detection (negative)

### Conventional markers of infectivity do not associate with aerosol production

Sputum was spontaneously produced by 18 of our 24 (75%) participants at 51 of a possible 192 (26.5%) time points with 1 to 8 samples produced and total volumes between 1 and 185 ml (Fig 2). Eight sputum samples from three patients were excluded due to processing errors. There was no observable diurnal pattern in either sputum production or Mtb content per sputum sample (Fig 3). While 24 hour Mtb sputum content was significantly higher than that in aerosol (p=0.02), in four participants the converse relationship was apparent (fig 2). We found no association between sputum and aerosol Mtb production either over the whole over 24 hours (p=0.06) or within the four patterns we have described (p=0.36). Interestingly, both the negative aerosol producers had detectable Mtb in sputum (fig 2).

The median cough frequency was 466 per 24 hour (IQR:234-551). Two participants (3&6) produced greater than 2000 coughs over the 24 hour sampling period, nearly 20-fold more than the lowest value of 39 (Figure 2). No association was seen between cough frequency and Mtb output detected in aerosols. This was investigated in three ways; between level of IS6110 detected in each hour of MASS and the corresponding cough frequency for that hour (p=0.27), between the 24 hour aerosol IS6110 output and the 24 hour cough frequency for each participant (p=0.63) and within each participant’s eight aerosol samples and cough frequency for each corresponding hour (Supplemental data).

Of the 33 aerosol samples that were negative for Mtb taken from 7 participants, 30 had coughs detected during sampling. The median cough frequency for these aerosol negative samples was 8 (2-37) which is similar to that demonstrated by the whole cohort (10 (2-22)). The discordance between cough frequency and MTb aerosol production was most evident in the 6 samples that recorded more than 50 coughs/hour (range 54-318).

Neither extent of disease nor presence of cavities on CXR were associated with total aerosol Mtb production (p=0.52). Sputum samples used in assessing participant eligibility, taken 24 hours prior to study commencement yielded AFB and Xpert grade as well as MGIT days to positivity, none of which showed correlated with 24 hour Mtb aerosol production. This can be seen in table 2 alongside other standard patient associated features (also not associated) and is equally true for the repeated analysis using RD9 normalised Mtb signals in aerosol and sputum.

**Table 2.** Predictors of PCR signal abundance in Sputum and Aerosol output over 24 hours. Data represents the ability of variables to predict levels of IS6110 and RD9 signals detected in Aerosol over 24 hours analysed by Spearman’s Correlation co-efficients for continuous variables and Mann Whitney U tests for categorical data. ^*^ CD4 recorded for all 20 HIV positive pts ^#^ CXR grade and presence of cavities for 23 pts ^¥^ Sputum AFB grade available for 17 pts ^$^ Sputum culture results available for 16 pts ^β^ VAS recorded for 22 pts.

| Predictors | 24 hour Aerosol IS6110 Output |  | 24 hour Aerosol RD9 Output |  |
| --- | --- | --- | --- | --- |
|  | Correlation coefficient (95% CI) | p-value | Correlation coefficient (95% CI) | p-value |
| Age | -0.26 (-0.61 - 0.17) | 0.21 | -0.13 (-0.52 - 0.30) | 0.53 |
| Gender |  | 0.89 |  | 0.47 |
| HIV status |  | 0.48 |  | 0.68 |
| CD4 count * | -0.28 (-0.65 - 0.20) | 0.23 | -0.11 (-0.53 - 0.36) | 0.65 |
| Duration of Symptoms (weeks) | 0.18 (-0.26 - 0.55) | 0.41 | 0.11 (-0.31 - 0.48) | 0.56 |
| CXR Grade<sup>#</sup> | -0.22 (-0.59 - 0.23) | 0.32 | -0.07 (-0.48 - 0.36) | 0.75 |
| Presence of cavitations on CXR<sup>#</sup> |  | 0.52 |  | 0.38 |
| Sputum AFB Grade<sup>¥</sup> | -0.08 (-0.54 - 0.41) | 0.75 | -0.02 (-0.49 - 0.47) | 0.95 |
| Sputum GeneXpert Grade | 0.02 (-0.40 - 0.43) | 0.92 | -0.04 (-0.45 - 0.37) | 0.84 |
| Sputum Culture<sup>§</sup> | 0.14 (-0.38 - 0.59) | 0.58 | 0.07 (-0.44 - 0.54) | 0.79 |
| Cough frequency | 0.10 (-0.33 - 0.50) | 0.63 | 0.07 (-0.35 - 0.47) | 0.73 |
| Patient perception of cough severity (VAS)<sup>β</sup> | 0.28 (-0.18 - 0.63) | 0.21 | 0.31 (-0.15 - 0.65) | 0.53 |
| Started TB medication |  | 0.68 |  | 0.91 |
| 24 hour Sputum PCR signal content | 0.17 (-0.30 - 0.57) | 0.47 | 0.15 (-0.31 - 0.56) | 0.49 |

Previous studies report associations between cough and AFB grade, extent of disease on CXR and days to culture positivity in liquid culture and this was confirmed in our cohort. Like Turner et al (ref) we did not find that cavitations and cough were correlated (p=0.88). (see supplemental table). Consistent with previous studies daytime cough frequency was higher than night time (p=0.04), women had a higher cough frequency than men (p=0.04) and cough frequency correlated with participants’ reported cough using a visual analogue scale (VAS) (ρ=0.50 (0.09 − 0.80) p=0.02).

### Potential Implications for TB Diagnosis

Mtb was detected by mask sampling at 166/192 (86.5%) possible sampling time points. In contrast, Mtb was detected from sputum in 38 of the 184 assessable (20.7%) sampling time points.

In the 43 sampling periods for which concomitant sputum and aerosol were available Mtb was detected by either aerosol or sputum in all sampling periods. However, there was discordance between the sampling methods for 12 of these periods. For 7 of these periods, sputum was Mtb positive and aerosol negative, while in the other 5 periods, aerosol identified Mtb but sputum was negative.

## Discussion

Using adapted face masks, we have been able to gain insight into the daily Mtb output in patient-generated aerosols. In contrast to established sampling methods, the MASS is simple, well tolerated and compatible with routine clinical practice. We have also presented evidence that conventionally applied indicators of individual infectivity do not predict aerosol output and that the MASS approach may have value as a broadly applicable non-invasive tool for both diagnosis and informing the likely infectivity of the active case. This study raises questions regarding current strategies widely used to assess patient infectivity.

While the work of Riley and Wells made it clear that aerosols were critical to transmission in guinea pig facilities^15-17^, only the work of Jones-Lopez and colleagues using the CASS has linked individual aerosol output levels with community transmission^3^. The work presented here, establishes the value of the MASS to recognise differences in aerosol production between and within individuals but not its link to transmission. It would none the less be surprising if the quantity of Mtb aerosols captured in MASS was not associated with transmission.

### Four patterns of aerosol production were observed and no diurnal pattern was evident

TB transmission is likely dependent on the number and quality of mycobacteria expelled into aerosol, however, until this study there has been no direct information on the pattern with which this occurs spontaneously. We identified four distinct groups of aerosol Mtb production over 24 hours. We were unable to find any features of their clinical presentation, particularly CXR that would explain these differences.

We found no evidence of a diurnal pattern in aerosol Mtb production. While this might reflect a bias of hospitalisation disrupting natural diurnal rhythm, a diurnal pattern of cough pattern was observed that is concordant with other studies^10,11,13^. It is also interesting that 53 positive aerosol samples were collected from participants that were directly observed to be sleeping for the full hour of sampling.

### Aerosol Production was not associated with traditional markers of infectivity

In the 43 sampling periods where both sputum and aerosol were available, Mtb content within these samples were not correlated. Moreover, the 24 hour output of bacilli in sputum did not predict that detected in aerosol and we conclude for individuals within this cohort, sputum content could not be used to predict aerosol output. This was also the case for Sputum AFB and Xpert MTB/RIF grade along with time to positivity in liquid culture. Recent data from Patterson and colleagues shows a similar discordance between sputum and aerosol production by participants in a South African setting^18^.

We observed no significant relationship between cough frequency and aerosolised Mtb production either for the full 24 hours of the study or at each individual time point assessed. In the Patterson study spontaneous cough frequency correlated with culturable bio-aerosol production detected in a specially designed cabinet^18^. Using the CASS approach, where patients were invited to cough, Fennelly and colleagues noted a trend associating cough frequency with colony forming units^2^ and more recently they reported an association with cough strength^3^. These groups, like us, failed to find any association between extent of disease and presence of cavitation on CXR with aerosol output. The reason for some of difference in results concerning cough may reflect the difference in sampling methods. The CASS and cough cabinet used by these groups selectively capture Mtb remotely from either a drum or large cabinet, in which the particles have time to desiccate into the respirable range. MASS, on the other hand, is non-selectively sampling both aerosols and droplets in a more proximal position.

### Relevance to Transmission

Comparing MASS with the CASS we note that output in the former reflects spontaneous expectoration over an hour whilst the later, a short 10 minute directed voluntary cough sample. It should also be appreciated that CASS captures colony forming bacilli assignable within the respirable range by use of the Andersen sampling system^2,3,19^. Through multiple sampling it has been possible to readily assess the consistency of output in our subjects, while no similar data is currently available for CASS.

It is striking that two highly variable aerosol producers were distinct in their 24 hour output from others in this HIV dominated group by more than an order of magnitude. This frequency concurs remarkably well with the findings of Escombe and colleagues in that 8.5% of an exclusively HIV cohort were highly infectious^20^. In considering how the patterns we have observed may relate to recognised individual events, we suggest that transmission from highly variable producers maybe associated with infection acquired from transient contact as a stochastic event, while that from the consistent producers may explain the utility of cumulative exposure time to inform linear risk in household and other close contacts. The relative importance of these two distinct mechanisms to transmission within populations remains uncertain.

In considering the relevance of sputum smear positivity to transmission, a recent meta-analysis, based on 37 studies, gave an adjusted odds ratio of 2.15 (1.47-3.17)^1^. Such population level studies contrast with a key CASS finding demonstrating a strong association (greater than 5 fold) between aerosol bacillary burden and transmission, while no such significant association was observed with sputum AFB grade. These authors also noted that ‘although aerosol production was correlated with sputum AFB smear grade in this and our previous studies, most subjects with high-grade smear (i.e AFB >3+) where aerosol negative’^3^. These findings accord with the dissociation between the aerosol and sputum results recorded here.

Considering reported cough, in several studies, various levels of association with transmission have been reported^1,3,12,18,21^. In the two studies where cough was independently assessed, a modest association with transmission was reported. Turner and colleagues using 24 hour data from the Leicester Cough Monitor, found an odds ratio of 1.30 (1.04-1.64)^12^ while Loudon and Spohn assessed night time cough counts and showed a trend to association with transmission (p=<0.20)^21^. In the latter regard, it is striking that we detected positive aerosols in 6 sleeping individuals where no cough was recorded on 11 separate occasions. It is also interesting that there were 6 occasions with high cough frequency (>5× the median cough frequency for the whole cohort) and no aerosolised Mtb detected. Although this could suggest a difference in our study population, in every other respect cough was predicted by the same patient indices (sputum and CXR grade) as the previous studies^12,18,21,22^.

The meta-analysis of Melsew and colleagues report and adjusted odds ratio of 1.90 (1.26-2.84) for the relationship between caviatory lung disease and transmission^1^. As reported by Jones-Lopez et al^3^, we found no association between aerosol Mtb production and extent of disease or the presence of cavitations on CXR.

Traditional indices of sputum grade, cough frequency and CXR changes are widely applied in clinical and public health practice to assess infectivity at the individual level. It is therefore surprising that neither our study, nor the CASS studies, show an association between these factors and Mtb aerosol production. Given the strength of association of aerosol Mtb production with household transmission found by Jones-Lopez et al^3^ we consider, at the very least, the need to develop and adopt additional tools to support risk assessment for transmission.

### Potential diagnostic Implications

Although it was not our original aim to determine the diagnostic value of MASS, we find it notable that aerosol samples were successfully obtained in every sampling period from every individual while this was only the case for sputum in less than one third of occasions. Given that Mtb was detected in 86.5% of aerosol samples and 88.4% of sputum samples, we suggest that aerosol sampling may offer a comparable result more frequently. Considering the ways in which mask sampling could be incorporated into clinical practice we note that this would be particularly advantageous in those patients that do not produce sputum or are smear negative at the first clinical encounter. Incorporating this approach into the primary diagnostic pathway could contribute to identifying the ‘missing millions’ identified by the WHO^23^.

### Limitations

To our knowledge this is the first observational study in which spontaneous output of Mtb in sputum and aerosol has been assessed over 24 hours.

The general applicability of our findings must be reviewed with caution. Firstly, the continuous monitoring of patients in this study required selection of hospitalised patients; this meant our participants tended to have more complicated or advanced disease. Secondly, the rate of HIV-TB co-infection in our cohort was higher than the WHO’s estimated rate of 57% in South Africa in 2015^24^. Thirdly, by restricting our eligibility criteria to sputum AFB/Xpert MTB/RIF positivity, our results may not apply to TB patients who are sputum negative or unproductive of sputum. Finally the small sample size of this study may have influenced our findings and made drawing conclusions about predictors of aerosol production within the groups difficult. That said, our findings do provide synergy with other groups with respect to some aspects of sputum bacillary load, cough and chest xray findings allow some confidence in its generalisability.

The choice of IS6110 as a multi-copy gene target, whilst providing sensitivity, complicates comparisons between patients within our cohort. Our normalisation to RD9 gives us some confidence in the comparisons we can draw. Two samples analysed for RD9 and IS6110 were RD9 negative but IS6110 positive supporting the choice of IS6110 for the majority of the analysis. We cannot say what proportion of the signals detected here represent potentially infective bacteria as our PCR would have detected DNA from both alive and dead bacteria. We note that signals from viable bacteria comparable to those detected here, were reported in our previous study^4^. Moreover, the culturable signals noted above in association with cough^2,3,19^ would not have detected in the bacterial that are incapable of forming colonies on solid media and require liquid culture and supplementation for their growth^25^. Although our group has previously demonstrated high numbers of Mtb with these requirements in sputum^25^, their occurrence in aerosol is has yet to be determined.

## Conclusion

We conclude that our primary hypothesis that traditional markers of infectivity should determine aerosol production, is not supported by our findings. Whether this reflects weakness in these markers or deficiencies in aspects of our study design will require further investigation.

## Acknowledgements

The authors acknowledge the invaluable contribution made by Ms. Sherrie Van Zyl, Ms Helen Sithole and Professor D. Van Zyl along with all the clinical and nursing staff at Kalafong, Tshwane District and Steve Biko Academic Hospitals in Pretoria for assistance with study recruitment. They also thank Professor Ed Nardell for support and study critique.

## Supplemental Data

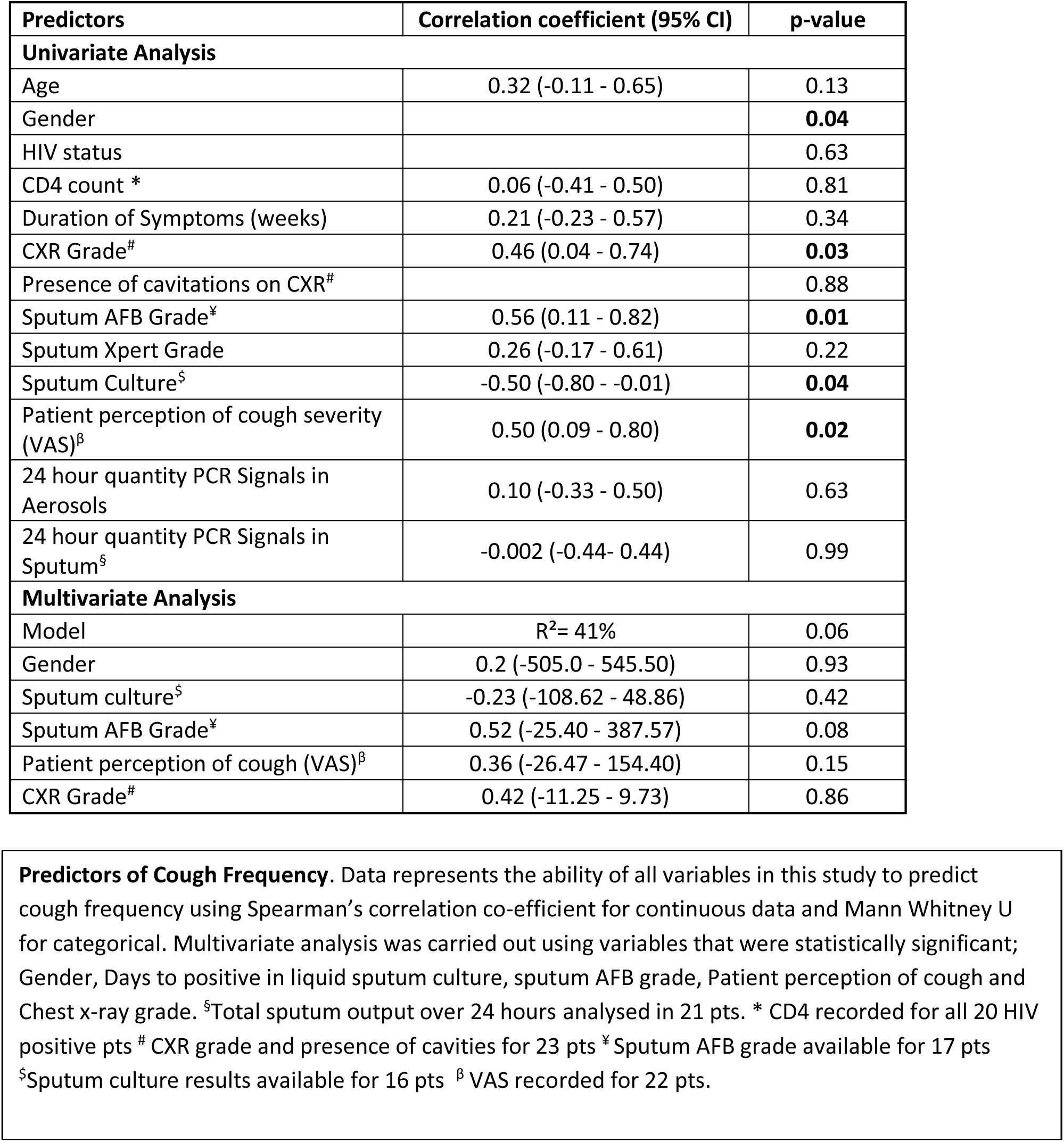

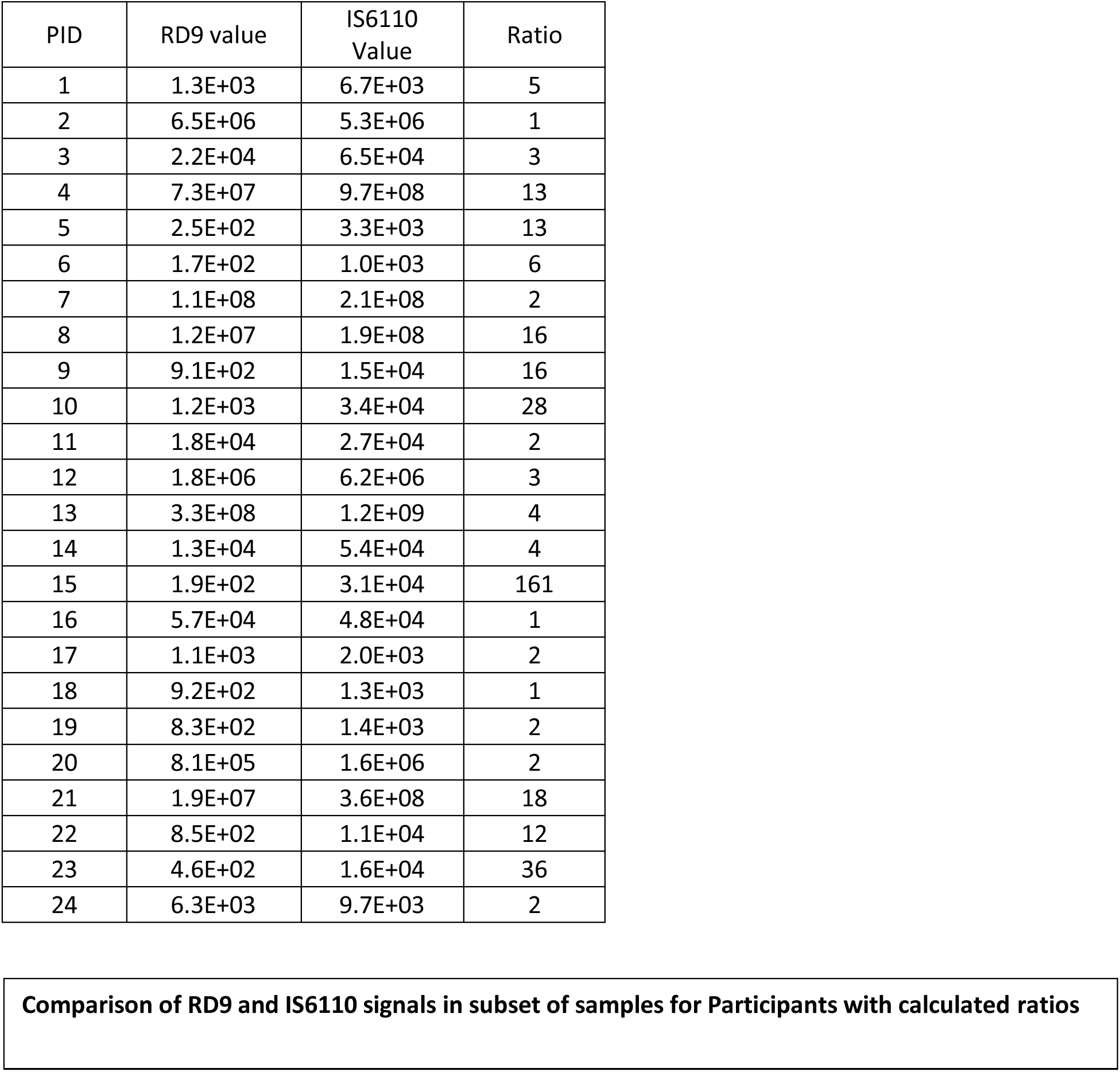

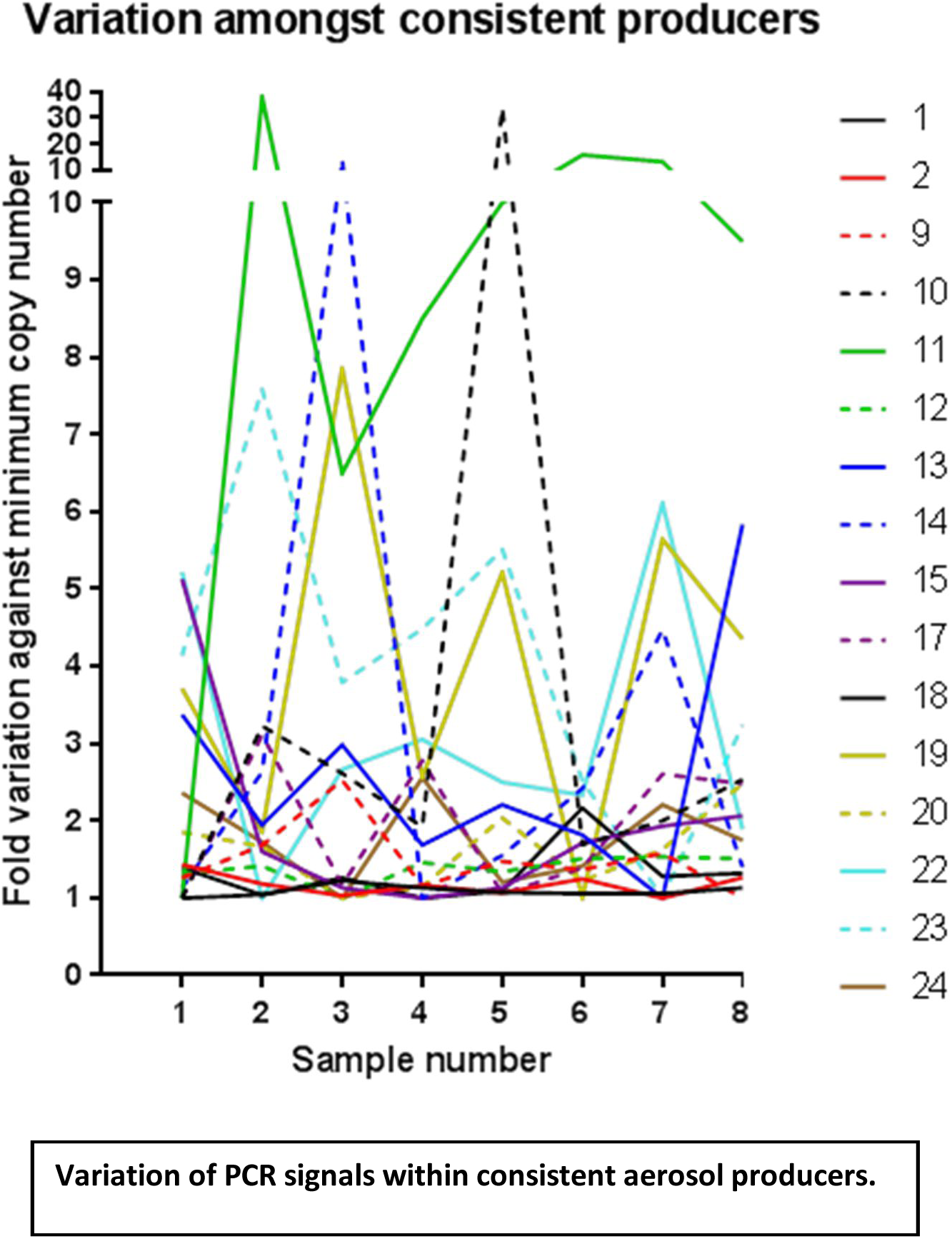

### Limit of Detection Methods

Mid – exponential Mtb H37Rv underwent serial 10-fold dilution to 10^7^ and filters were contaminated with 100μl of each dilution in 10μl drops across the surface of the filter. Mycobacterial DNA was isolated and extracted before IS6110 copies were quantified. A negative control arm was analysed using un-contaminated filters. This experiment was conducted in technical triplicate. The CFU of original Mtb suspension was calculated using the drop plate method described by Hoben and colleagues^1^. The number of Mtb genomes recovered was calculated by dividing the absolute quantification of IS6110 by 16 (IS6110 copy number in H37Rv).

A limit of detection (LoD) for this method was calculated using the following formulae outlined by Armbruster and colleagues^2^, the results from the dilution series and a further 9 blank filters which were processed in the same way in order to calculate the limit of the blank (LoB).

LoB = meanBlank + 1.645(SD_Blank_)

LoD = LoB + 1.645(SD_low concentration sample_).

Quantification of Mtb genomes recovered from differing dilutions of contaminated filters using the NaOH and in house method is displayed in Figure 6. The mean (SD) of the IS6110 copies recovered from the 12 bank filters processed was calculated as 218.9(27.2) and so the LoB was calculated as 264. The standard deviation of the lowest dilution that was above the blank (i.e. 1.8×10^3^) was calculated as 162.5.

Therefore LoD was calculated as:

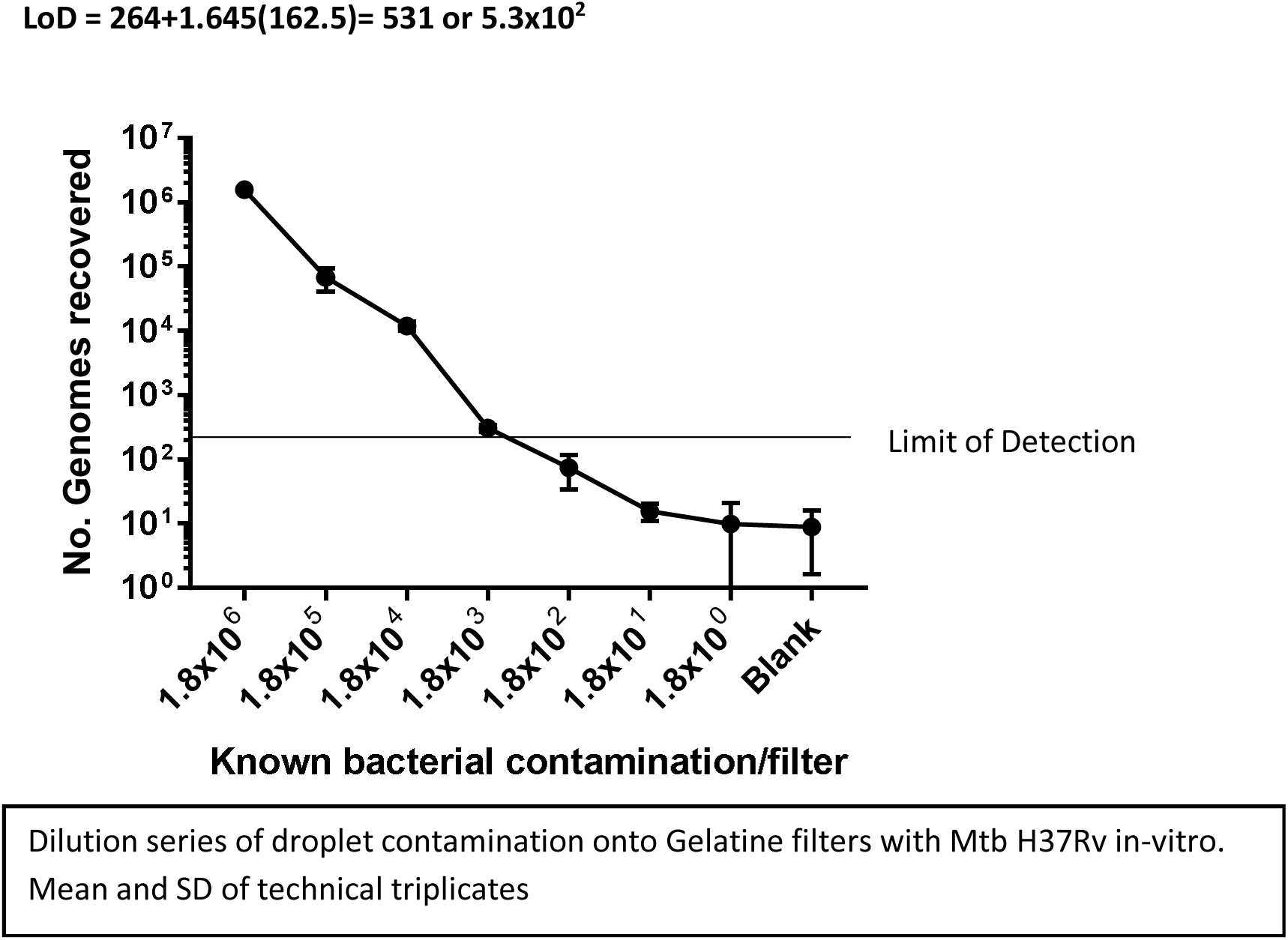

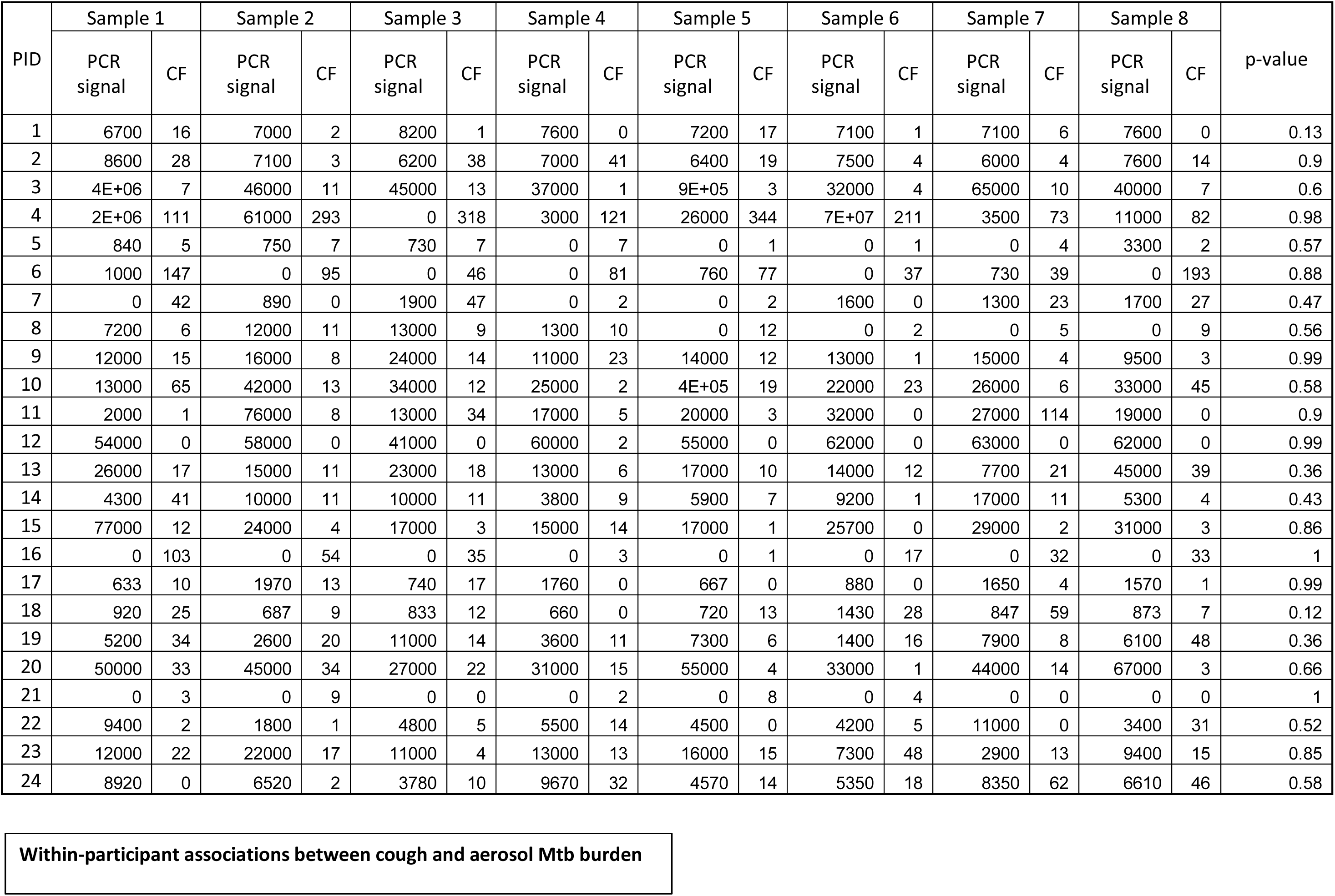

1. Hoben H, Somasegaran PJA, microbiology e. Comparison of the pour, spread, and drop plate methods for enumeration of Rhizobium spp. in inoculants made from presterilized peat. 1982; **44**(5): 1246–7.
2. Armbruster DA, Pry T. Limit of blank, limit of detection and limit of quantitation. *Clin Biochem Rev* 2008; **29 Suppl 1**(Suppl 1): S49–52.

## References

1. Melsew YA, Doan TN, Gambhir M, Cheng AC, McBryde E, Trauer JM. Risk factors for infectiousness of patients with tuberculosis: a systematic review and meta-analysis. Epidemiology and infection 2018; 146(3): 345–53.

2. Fennelly KP, Martyny JW, Fulton KE, Orme IM, Cave DM, Heifets LB. Cough-generated aerosols of Mycobacterium tuberculosis: a new method to study infectiousness. Am J Respir Crit Care Med 2004; 169(5): 604–9.

3. Jones-Lopez EC, Namugga O, Mumbowa F, et al. Cough aerosols of Mycobacterium tuberculosis predict new infection: a household contact study. Am J Respir Crit Care Med 2013; 187(9): 1007–15.

4. Williams CM, Cheah ES, Malkin J, et al. Face mask sampling for the detection of Mycobacterium tuberculosis in expelled aerosols. PLoS One 2014; 9(8): e104921.

5. Turapov O, O’Connor BD, Sarybaeva AA, et al. Phenotypically Adapted Mycobacterium tuberculosis Populations from Sputum Are Tolerant to First-Line Drugs. Antimicrob Agents Chemother 2016; 60(4): 2476–83.

6. C.A. Reddy TJB, J.A. Breznak, G.A. Marzluf, T.M. Schmidt, L.R. Snyder. Methods for General and Molecular Microbiology. Third ed: ASM Press; 2007.

7. Akkerman OW, van der Werf TS, de Boer M, et al. Comparison of 14 molecular assays for detection of Mycobacterium tuberculosis complex in bronchoalveolar lavage fluid. J Clin Microbiol 2013; 51(11): 3505–11.

8. Dorak MT. Real Time PCR Taylor & Francis Group; 2006.

9. Chae H, Han SJ, Kim SY, et al. Development of a one-step multiplex PCR assay for differential detection of major Mycobacterium species. 2017: JCM. 00549–17.

10. Birring SS, Fleming T, Matos S, Raj AA, Evans DH, Pavord ID. The Leicester Cough Monitor: preliminary validation of an automated cough detection system in chronic cough. Eur Respir J 2008; 31(5): 1013–8.

11. Sinha A, Lee KK, Rafferty GF, et al. Predictors of objective cough frequency in pulmonary sarcoidosis. Eur Respir J 2016; 47(5): 1461–71.

12. Turner RD, Birring SS, Darmalingam M, et al. Daily cough frequency in tuberculosis and association with household infection. Int J Tuberc Lung Dis 2018; 22(8): 863–70.

13. Chan KK, Ing AJ, Laks L, Cossa G, Rogers P, Birring SS. Chronic cough in patients with sleep-disordered breathing. Eur Respir J 2010; 35(2): 368–72.

14. Ralph AP, Ardian M, Wiguna A, et al. A simple, valid, numerical score for grading chest x-ray severity in adult smear-positive pulmonary tuberculosis. Thorax 2010; 65(10): 863–9.

15. Riley R, Mills C, Nyka W, et al. Aerial dissemination of pulmonary tuberculosis. A two-year study of contagion in a tuberculosis ward. 1959; 70(2): 185–96.

16. Riley R, Mills C, O’grady F, Sultan L, Wittstadt F, Shivpuri DJARoRD. Infectiousness of air from a tuberculosis ward: ultraviolet irradiation of infected air: comparative infectiousness of different patients. 1962; 85(4): 511–25.

17. Sultan L, Nyka W, Mills C, O’grady F, Wells W, Riley RJARoRD. Tuberculosis disseminators: a study of the variability of aerial infectivity of tuberculous patients. 1960; 82(3): 358–69.

18. Patterson B, Morrow C, Singh V, et al. Detection of Mycobacterium tuberculosis bacilli in bio-aerosols from untreated TB patients. Gates Open Res 2017; 1: 11.

19. Jones-Lopez EC, Acuna-Villaorduna C, Ssebidandi M, et al. Cough Aerosols of Mycobacterium tuberculosis in the Prediction of Incident Tuberculosis Disease in Household Contacts. Clin Infect Dis 2016; 63(1): 10–20.

20. Escombe AR, Moore DA, Gilman RH, et al. The infectiousness of tuberculosis patients coinfected with HIV. PLoS Med 2008; 5(9): e188.

21. Loudon RG, Spohn SK. Cough frequency and infectivity in patients with pulmonary tuberculosis. Am Rev Respir Dis 1969; 99(1): 109–11.

22. Proaño A, Bravard MA, López JW, et al. Dynamics of cough frequency in adults undergoing treatment for pulmonary tuberculosis. 2017; 64(9): 1174–81.

23. Organization WH. Global tuberculosis report 2017.. 2018.

24. Organization WH. Global tuberculosis report 2016. 2016.

25. Mukamolova GV, Turapov O, Malkin J, Woltmann G, Barer MR. Resuscitation-promoting factors reveal an occult population of tubercle Bacilli in Sputum. Am J Respir Crit Care Med 2010; 181(2): 174–80.

